# LEM domain proteins control the efficiency of adaptation through copy number variation

**DOI:** 10.1101/451583

**Authors:** Paolo Colombi, Diane E. King, Jessica F. Williams, C. Patrick Lusk, Megan C. King

## Abstract

While loss of genome integrity is at the basis of numerous pathologies, including cancer, genome plasticity is necessary to adapt to a changing environment and thus is essential for long-term organismal success. Here we present data supporting a targeted mechanism that promotes adaptation to environmental stress by driving site-specific genome instability tied to transcriptional induction and the formation of RNA-DNA hybrids. Using an *in vitro* evolution assay we observe that the inner nuclear membrane LEM domain proteins Heh1 and Heh2 play antagonistic roles in inhibiting or promoting adaptation through copy number expansion, respectively, which is also reflected in their genetic interaction networks with genes responsible for transcription-dependent genome instability. Taken together, our data suggest the existence of a LEM domain protein-mediated mechanism by which an immediate transcriptional response to a changing environment drives targeted genome instability to promote increased variation on which selection can act to support long-term adaptation.

## Introduction

The success of an organism relies in part on its ability to maintain fitness in the face of a changing environment, often sensed by increased cellular stress. It has been proposed that multiple mechanisms drive adaptation, differing in the time scale over which they take place, how long they can be sustained, their heritability, and their reversibility(Yona et al., 2015). These mechanisms fall into two major classes: non-genetic changes rooted in physiological responses that drive transcriptional and/or epigenetic modifications, and genetic changes occurring through point mutations and/or gene copy number variation (CNV). A model in which rapid, non-genetic physiological responses could drive adaptive genetic changes has been suggested to couple these two mechanisms [1]. Consistent with this model, recent work suggests that genes poised to respond transcriptionally to environmental challenges may also be subject to “stimulated” adaptive copy number variation of these genes [2]. However, the mechanisms coupling transcriptional up-regulation to local genome instability that can drive copy number variation have yet to be fully delineated.

In neo-Darwinian theory, genetic change occurs gradually and constantly in a stochastic fashion independent of the environment, becoming fixed in the genome as a consequence of providing a fitness advantage. However, in principle an inducible system where bursts of mutations are produced only in conditions of stress [3, 4] provides advantages over a system based on a constitutive, high mutator phenotype that could prove deleterious under conditions where an organism is well-adapted [5]. Indeed, Barbara McClintock provided the first evidence that elevated insertion-deletion (indel) events, CNV, and other genome rearrangements are driven by elevated transposon activity in response to stress, which acts independently of stochastic environmental or metabolic damage to the DNA [6]. A similar phenomenon has been observed in prokaryotes, in which a mutagenic “SOS response” and DNA break repair mechanisms are up-regulated in response to stress (reviewed in [7]. Interestingly, the role of “stress induced mutagenesis” (SIM) might occur in diverse eukaryotes from yeasts [8-10], to algae [11], nematodes [12], and human cancer cells [13, 14], suggesting its broad conservation. However, as the SOS response appears to be prokaryote-specific, we lack a coherent framework that explains the mechanistic details of how stress may drive mutagenic processes in eukaryotic organisms.

Physiological changes including transcriptional induction and the landscape of epigenetic modifications that facilitate the rapid response to stress have also been suggested to drive genome instability [15, 16], thereby coupling non-genetic and genetic sources of adaptive potential, although whether high levels of transcription drive an increase in mutation rate in the absence of selection remains debated [17-19]. The ribosomal DNA repeats in *S. cerevisiae* provide an example of a genomic locus undergoing controlled genetic changes in a transcription-dependent manner that is linked to epigenetic modifications [20-24]. In this case, the repetitive nature of the rDNA supports frequent CNV (also called “repeat instability”); likely fitness advantages provide a selective pressure for maintaining an ideal rDNA copy number, for example cultures of strains engineered to have decreased rDNA copy number rapidly “acquire” additional rDNA copies [25]. Hints that an active mechanism may explain such observations have emerged. For example, it has been suggested that cells possess mechanisms to control rDNA copy number by modulating either the frequency of local DNA stability within the locus and/or the mechanism of homology-directed repair (HDR) used to repair local DNA lesions [26, 27]. One influence on this process may be the sub-nuclear compartmentalization of the rDNA in the nucleolus, which is associated with the inner nuclear membrane (INM) in yeasts. Indeed, disrupting an rDNA tether to the nuclear envelope, provided by the integral inner nuclear membrane LEM (LAP2, emerin, MAN1) domain protein Heh1/Src1, increases rDNA CNV [28, 29], although the mechanism remains obscure. Moreover, it remains largely unexplored if similar mechanisms act elsewhere in the genome to modulate the frequency of CNV or point mutations, and, if so, how such mechanisms might contribute to adaptation.

Here we identify a stress-response pathway that modulates cellular adaptation through CNV. Interestingly, the extent of CNV is controlled at the INM, where genetic deletion of *HEH1* drives increased CNV through a mechanism that requires its ohnologue, *HEH2*[30]. Our findings support a model in which Heh2 acts antagonistically to Heh1 through a pathway that leads to the transcription-dependent formation of RNA-DNA hybrids. We suggest a model in which non-genetic changes at specific genetic loci that associate with the nuclear periphery drive local CNV to promote adaptation.

## Results

To study the innate mechanisms by which cells adapt to changes in their environment through CNV, we adopted the *S. cerevisiae* multicopy *ENA* gene locus as an experimental model, which comparative genomics approaches had suggested to undergo high levels of CNV [31]. Genome assemblies of budding yeast strains suggest that up to five *ENA* gene copies (with ≈98/99% identity) reside in a tandem array[32-35]; our WT W303 strain has four tandem *ENA* coding sequences, as confirmed by PCR, which we refer to here as *ENA1-4* (Figure 1A; Supplemental Figure 1A-B). Consistent with the idea that the *ENA* locus encodes pumps responsible for salt efflux, it is rapidly induced ≈3-fold in the presence of 100 mM LiCl (Figure 1B) and is essential under high salt conditions [36, 37] (Figure 1C).

**Figure 1.**
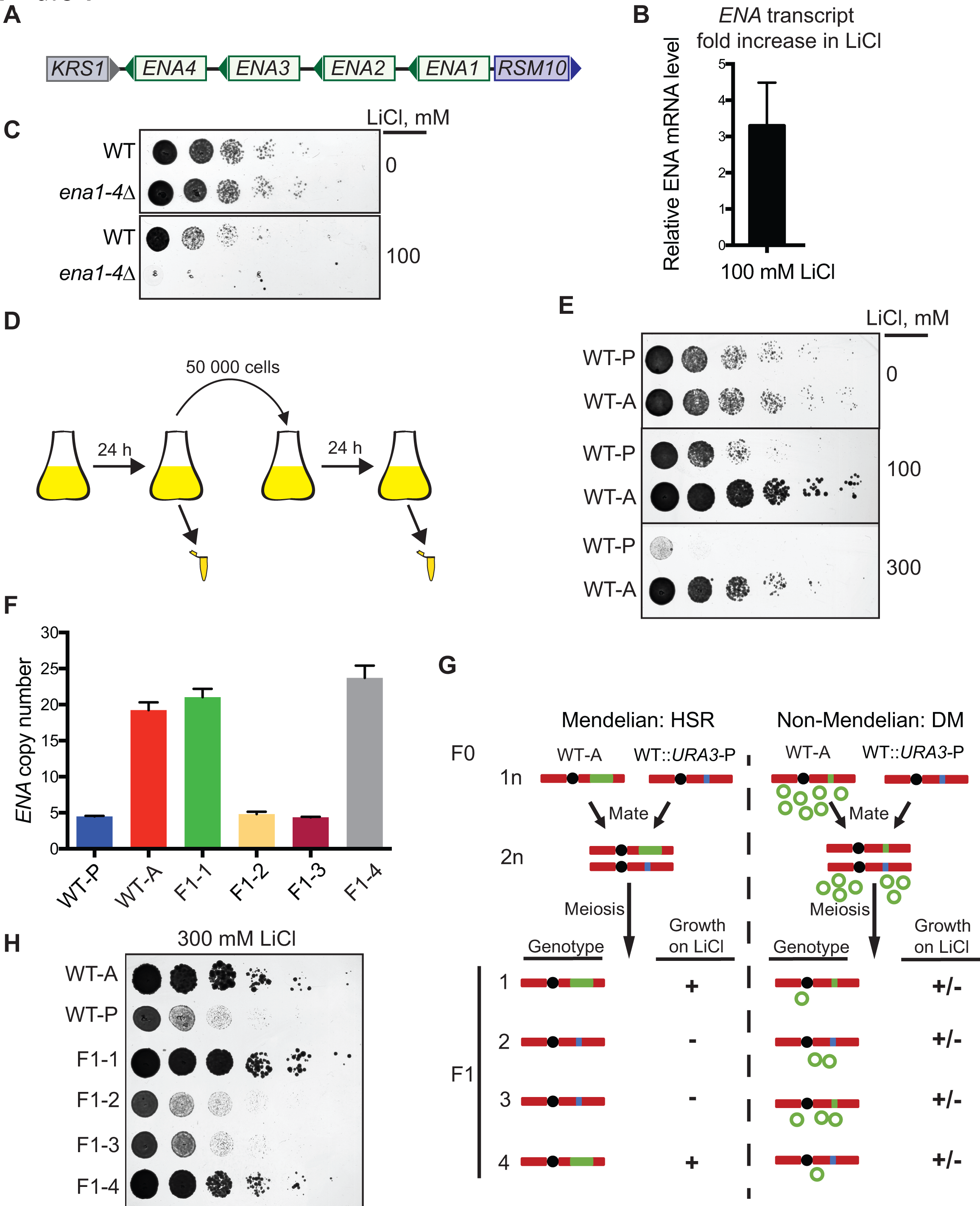
*ENA* copy number expansion favors adaptation to high salt. (A) Schematic of the *ENA* genomic locus, including the *ENA* genes (green, numbered from 1 to 4), *RSM10* (upstream of the *ENA* locus, violet), and *KRS1* (downstream of the *ENA* locus, grey). Arrowheads indicate the direction of transcription. (B) The level of the *ENA1-4* transcript increases after 10 min. in the presence of 100 mM LiCl. RT-qPCR analysis of *ENA1*-*4* transcript in 100 mM LiCl normalized to 0 mM LiCl. Mean ± SD from 3 independent experiments. (C) Deletion of *ENA1*-*4* leads to cell death in the presence of 100 mM LiCl. Serial dilutions, 1:10, were grown on YPD plates with or without 100 mM LiCl. (D) Schematic of the *in vitro* evolution experiment. Cells were cultured for ≈200 generations in YPD containing 100 mM LiCl; every 24 h cells were diluted to 1,000 cells/ml in fresh media. (E) The adapted strain (WT-A) has a fitness advantage over the parental strain (WT-P) when grown on YPD plates containing 100 mM and 300 mM LiCl. Serial dilutions as in (C). Note that plates containing LiCl were imaged after growth for an additional 24 h. (F) The copy number of the *ENA* locus is increased in WT-A and its meiotic progeny (described in G-H) as measured by qPCR. Mean ± SD of three independent experiments. (G) Schematic of the predicted genotypic and phenotypic (relative growth in LiCl denoted by + and -) differences of progeny (F1) derived through Mendelian inheritance (homogenous staining region/HSR; expansion on the chromosome) and non-Mendelian double minutes (DM; generation of episomes) during meiotic segregation. The chromosome (in red) can contain the parental *ENA* genomic locus (in blue), or the adapted *ENA* locus (in green, HSR as single band or DM as circles). (H) The spores inheriting the adapted ENA locus (F1-1 and F1-4) retain improved growth on media containing 300 mM LiCl. Serial dilutions as in (C).

To identify pathways that impact *ENA* CNV, we devised an *in vitro* evolution assay wherein cells are grown in liquid medium containing 100 mM LiCl by serial culturing for ≈200 generations (see Materials and Methods)(Figure 1D). Prolonged culturing in LiCl resulted in WT adapted (“WT-A”) strains that tolerate high concentrations (up to 300 mM) of LiCl compared to the WT parental (“WT-P”) strain (Figure 1E). To test the underlying basis for improved fitness of WT-A, we measured *ENA* gene copy number by qPCR. The adapted strain expanded its *ENA* gene dosage by 500%, possessing an average of 20 copies per genome (Figure 1F). To differentiate whether the increase in gene copy number occurred through intrachromosomal or extra-chromosomal expansion (historically referred to as a “homogenous staining region” (HSR) or double minutes (DMs), respectively), we tested whether salt tolerance was inherited through a Mendelian or random genetic segregation in meiosis (Figure 1G). The WT-A strains was crossed with a WT strain with a genetic marker integrated adjacent to the *ENA* locus to monitor segregation; WT::*URA3*-P) and meiosis was induced. We observed a Mendelian inheritance pattern where two of the four progeny (F1 and F4) maintained ≈20 copies of the *ENA* locus (Figure 1F) with concomitant salt tolerance (Figure 1H). These data support the conclusion that cells adapt to high salt through an intrachromosomal expansion of the *ENA* genomic locus (≈20 *ENA* genes; Figure 1F-H), and not through the production of extrachromosomal DMs.

Copy number expansion is typically driven by non-conservative homology-directed repair processes downstream of replication fork collapse or the formation of DNA double-strand breaks[38]. As it has been suggested that high levels of transcription can drive local genome instability [15], and *ENA* expression is transcriptionally induced upon growth in high salt (Figure 1B), we next tested if transcription drives *ENA* locus instability. We devised a genome stability assay similar to that used to reveal the nonuniform rate of mutation across the budding yeast genome[39, 40]. Here, counter-selection for Ura3 activity, which converts 5-Fluoroorotic acid (5-FOA) to the toxic metabolite, 5-fluorouracil, allows the frequency of mutations in (or loss of) *URA3* to be defined. We inserted the *URA3* gene in three different positions at the *ENA* locus (Figure 2A) and measured the spontaneous instability at each position over a single cell cycle (2 hours growth before plating on 5-FOA; loss of *URA3* must occur prior to plating). In all positions, Ura3 activity is lost at rates at least an order of magnitude higher than at the endogenous *URA3* locus (≈10^-8^; Figure 2B), suggesting a high level of local instability. As expected due to flanking tandem repeats [41-44], the highest level of instability (≈10^-5^) is observed when *URA3* is inserted within the *ENA* copies (Figure 2B (“In”); in this case, the *URA3* gene is always lost, presumably by non-allelic homologous recombination; Supplemental Figure 2A). Interestingly, the instability downstream (3’) of the *ENA* gene locus (≈10^-6^) is markedly higher than upstream (5’; ≈10^-7^), suggesting a potential role for *ENA* transcription in modulating *URA3* stability. Consistent with this idea, inducing *ENA* gene expression by addition of LiCl during the single generation (2 hours) of release (from –uracil to 5-FOA) drives a higher frequency of 5-FOA resistant colonies specifically at *URA3* inserted downstream of the *ENA* locus (≈30%). Consistent with the idea that this effect is a direct result of transcription, it was abolished in cells harboring a loss of function allele of RNA polymerase II, *rpb1-1*[45, 46] (Figure 2C). In contrast, the region upstream of the *ENA* locus showed a lower frequency of 5-FOA resistant colonies in the presence of LiCl (≈40%) while the stability of the endogenous *URA3* locus is unaffected (Figure 2C).

**Figure 2.**
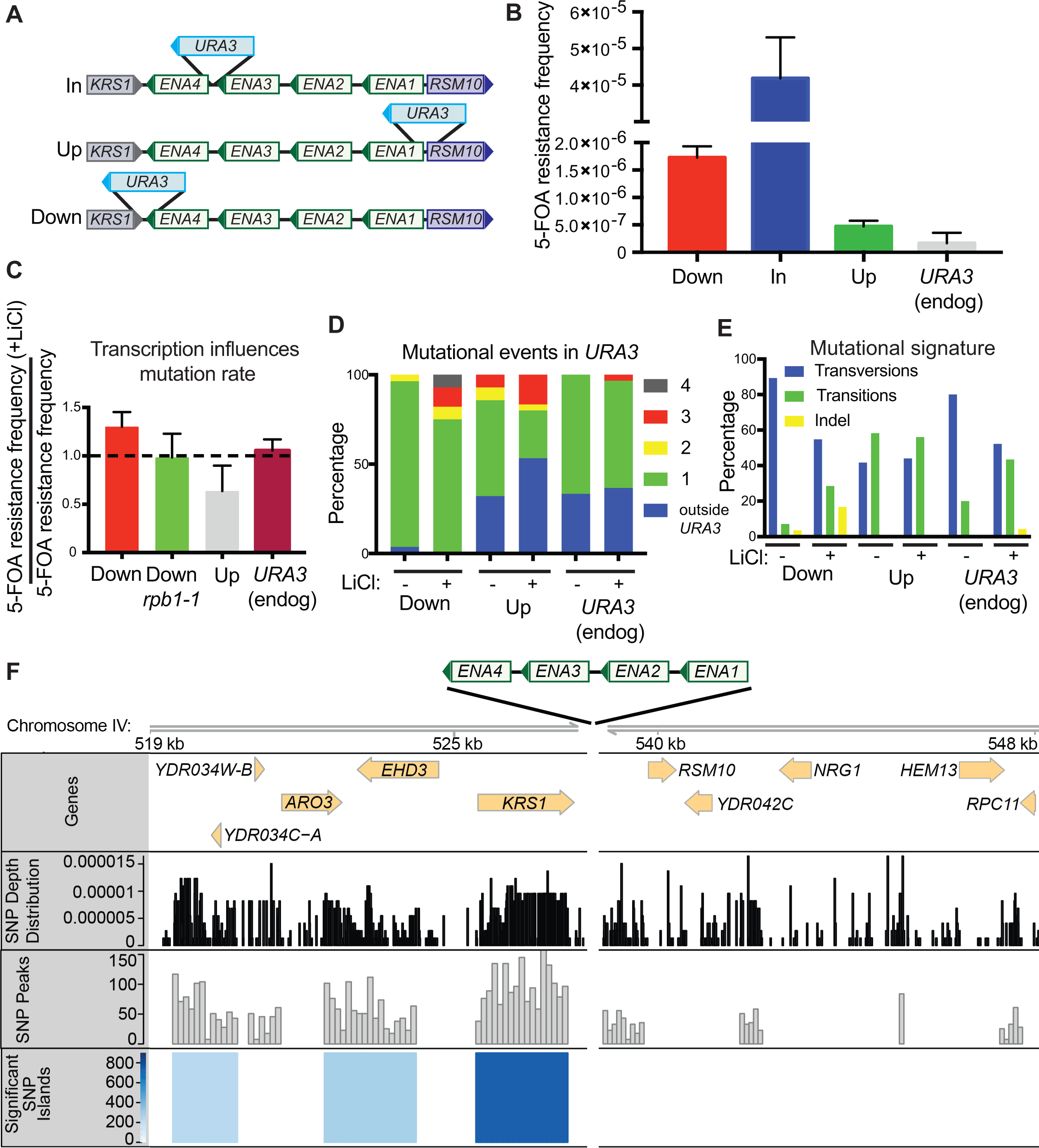
*ENA* locus instability is biased toward the 3’ region and is promoted by transcription. (A) Schematic representing the location of *URA3* insertions at the *ENA* locus. (B) The *ENA* genomic locus and its flanking regions have higher spontaneous instability compared to *URA3* at its endogenous locus. Measure of the loss of *URA3* activity as the rate of obtaining clones resistant to 5-FOA compared to total cells plated. Mean ± SD of three independent experiments. (C) Addition of LiCl to induce *ENA* expression (Fig. 1B) drives increased instability specifically downstream of the *ENA* locus. This effect is abrogated in cells harboring a mutation that ablates Pol II function (*rpb1-1*). Data is expressed as the ratio for LiCl-treated to untreated cells. Mean ± SD of three independent experiments. (D) Increasing *ENA* gene expression affects the number of mutational events within *URA3* specifically downstream of the *ENA* locus. In blue are events driving 5-FOA resistance outside the *URA3* gene. (E) Addition of LiCl to induce *ENA* expression alters the mutational signature specifically downstream of the *ENA* locus. (F) The SNP load is biased downstream of the *ENA* locus. SNP depth derived from 21 *S. cerevisiae* strains (black) processed to identify SNP peaks (gray), which were further analyzed to identify significant SNP islands (shaded in blue tones by score; see Methods).

5-FOA resistant colonies derived from strains with *URA3* inserted downstream of the *ENA* locus almost all map to the *URA3* gene, with an increase in the number of mutations per *URA3* gene upon exposure to LiCl (from ≈5% to ≈25% of 5-FOA-resistant clones have 2-4 mutations in *URA3*; Figure 2D). By contrast, the mutation profile of the upstream position is very similar to that of the endogenous locus, where ~30% of the 5-FOA-resistant colonies occur outside the *URA3* gene (Figure 2D) and does not change in response to LiCl. A more detailed mutational signature analysis shows that mutations driving 5-FOA-resistance at *URA3* integrated downstream of *ENA* are enriched in transversions (≈90%), which shifts to an increase of transitions (≈30%) and indels (≈15%) after LiCl exposure (Figure 2E); this contrasts with *URA3* integrated upstream of the *ENA* locus, which is unaffected by addition of LiCl and matches the transition/transversion rate typical of the budding yeast genome as a whole (≈0.6)[47]. At the endogenous *URA3* locus in the absence of LiCl we observe a mutational profile in line with previous studies [39](Figure 2E). Here we also observe a shift from transversions to transitions after LiCl treatment, although these events remain relatively rare given the low rate of recovering mutations at *URA3*(Figure 2B,D).

These observations suggest that local factors might drive genome instability and/or the fidelity of repair [40, 48]. To gain further insight we turned to a genome-wide analysis of single nucleotide polymorphism (SNP) distribution across twenty-one *S. cerevisiae* strains [49, 50], in which we defined peaks of significantly locally high SNP density (“islands”; Figure 2F; see Methods). Ranking SNP islands according to their integrated peak height and width to give rise to a SNP island score (see Methods), we found that the highest ranked islands (18 of the top 20; 41 of the top 50, Supplemental Figure 2B, Table S1) harbor genes that fall into at least one of two categories: subtelomeric genes and genes with paralogues (many of which are ohnologues arising from the whole genome duplication in the *S. cerevisiae* lineage), both of which are established to undergo rapid changes in sequence content [51-53]; indeed, none of the genes located in the top 20 SNP islands are essential. Surprisingly, the essential *KRS1* gene, which lies immediately downstream of the *ENA* locus, scored 21^st^ of all ranked islands (Table S1); the unusually high mutational frequency within *KRS1* was in fact noted in the initial comparative genomic sequence analysis of budding yeast strains [54]. More generally, the entire genomic region downstream of the *ENA* locus has a high SNP load compared to the region upstream of the *ENA* genes (Figure 2F), consistent with the heightened instability of *URA3* at this position.

To gain insight into the factors that might underlie the high SNP load and relative instability of the *KRS1*-*ENA* region, we investigated its sub-nuclear localization by integrating a lac Operator array just upstream of *ENA1* and monitored its position in cells expressing GFP-lacI and Hmg1-mCherry, a nuclear envelope/ER marker. Interestingly, the *ENA* locus is more strongly enriched at the nuclear envelope compared to the *URA3* locus (Supplemental Figure 3), raising the possibility that the nuclear periphery might play a role in modulating the stability of this genomic region. As Heh1 was previously implicated in modulating the stability of the rDNA [29], we investigated if Heh1 (and/or its ohnologue, Heh2; Figure 3A) influences the chromosomal expansion of the *ENA* locus in response to salt stress by carrying out the *in vitro* LiCl evolution experiment for WT, *heh1*Δ, *heh2Δ*, and *heh1*Δ*heh2*Δ strains. All four genetic backgrounds acquired improved fitness on media containing LiCl (100 mM and 300 mM) after 200 generations of culturing in 100 mM LiCl (Figure 3B). However, we uncovered marked differences in *ENA* copy number in the adapted strains by qPCR depending on genotype. In WT strains, *ENA* copy number is stable at 4 copies over 60 generations but increases to ≈6 at around 90 generations, and ultimately reaches ≈20 by 200 generations (Figure 3C). Interestingly, the *heh1*Δ strain undergoes a more rapid expansion of *ENA* copies, doubling to ≈8 at 60 generations and reaching a maximum copy number of ≈32 by 90 generations, which is maintained until ≈200 generations. Surprisingly, the ablation of the paralogous *HEH2* has the opposite effect, delaying the *ENA* copy number expansion at all time points. Cells lacking both *HEH1* and *HEH2* behave nearly like WT, suggesting the possibility of antagonism between these two ohnologues on *ENA* locus copy number.

**Figure 3.**
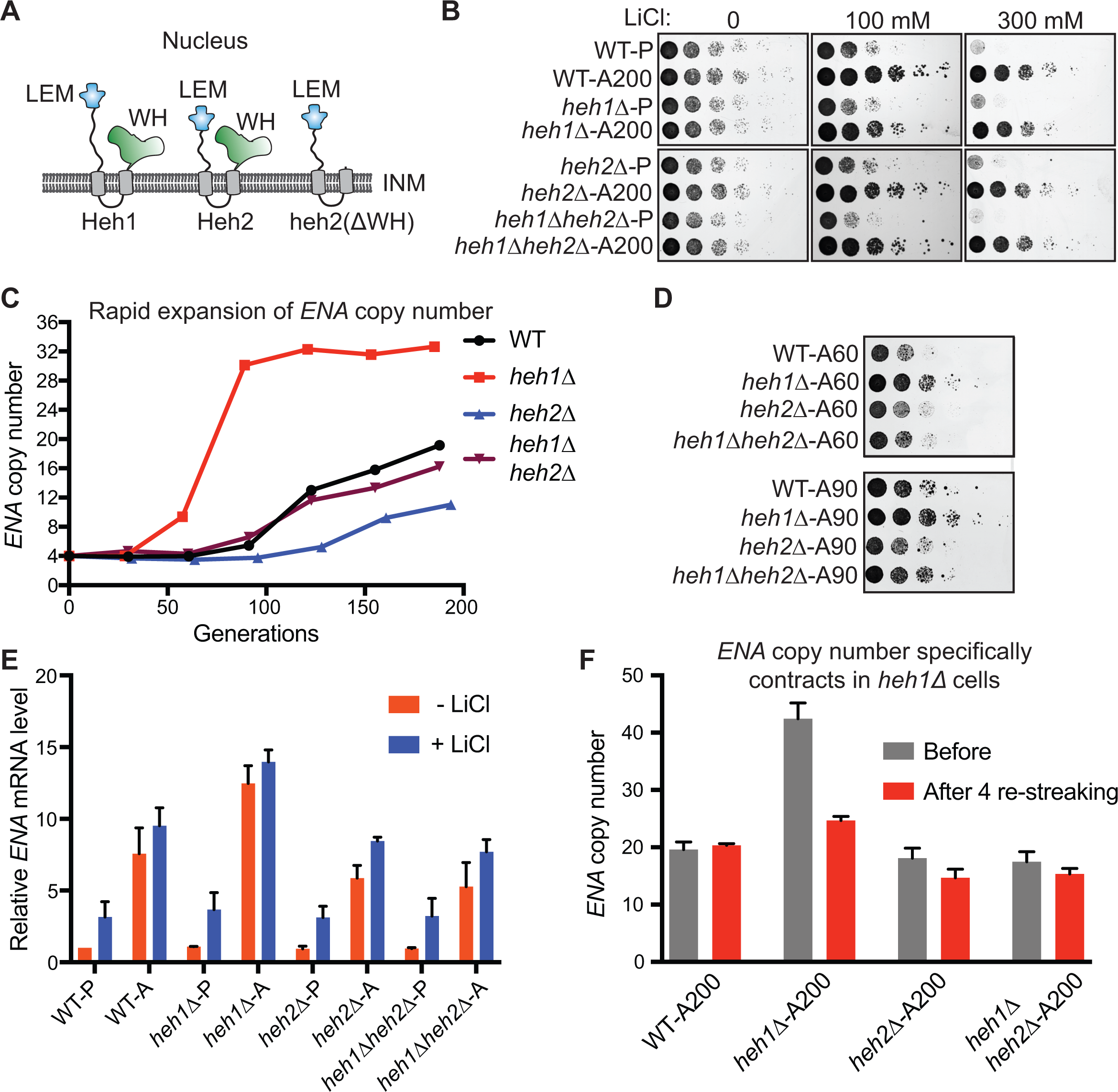
LEM domain proteins control adaptation efficiency through *ENA* copy number variation. (A) Schematic of the topology and domain architecture of Heh1 and Heh2. The conserved domains LAP2-emerin-MAN1 (LEM, blue), MAN1 C-terminal homology domain/winged helix (WH, green), and transmembrane domains (grey) are indicated. INM is inner nuclear membrane. (B) The adapted (A) strains have better fitness than the parental strains (P) growing on YPD plates containing 100 mM and 300 mM LiCl. The adapted strains were obtained after serial culturing (~ 200 generations) in YPD containing 100 mM LiCl. Serial dilutions as in Fig. 1C. (C) Deletion of *HEH1* and/or *HEH2* influences the rate of *ENA* copy number expansion. qPCR measures of the *ENA* copy number through the course of the *in vitro* evolution experiment. (D) Fitness comparison of the four genetic backgrounds at ~60 and ~90 generations. Serial dilutions as in Fig. 1C. (E) *ENA1-4* transcript levels increase in the adapted strains. Transcript levels of parental and adapted strains in control conditions (YPD) and after incubation for 10 min with 100 mM LiCl. The data are normalized to WT-P grown in YPD. Mean ± SD of three independent experiments. (F) *HEH1* influences the stability of the adapted *ENA* locus in the absence of selective pressure. qPCR analysis of *ENA* copy number of adapted strains before and after 4 serial re-streaking on media lacking LiCl.

The observed *ENA* copy numbers directly correlate with growth fitness in the presence of LiCl as only *heh1*Δ is resistant to growth in LiCl at ≈60 generations, while at ≈90 generations the *heh2*Δ is the least fit as it only has 4 copies (on average) of the *ENA* genes (Figure 3C-D). The copy number increase is driven by intra-chromosomal expansion as for WT strains in all cases (Supplemental Figure 4) and is reproducible (Supplemental Figure 5). Importantly, the influence of *HEH1* and *HEH2* on copy number expansion cannot be explained by changes in inherent salt tolerance or *ENA* expression at baseline as assessed by RT-qPCR (Figure 3B and E). Moreover, levels of the *ENA* transcript increase proportionally with gene copy number in the adapted strains, although the relative influence of LiCl on *ENA* transcript levels is diminished with increasing copy number (in the WT and *heh1*Δ backgrounds, compare parental to adapted strains, Figure 3E). Lastly, to gain insight into the stability of the expanded *ENA* copy number in the absence of salt stress, we serially re-streaked the adapted strains on rich media. Interestingly, only the *heh1*Δ strain shows a loss of *ENA* copy number (almost 50%), while the other strain backgrounds remain stable (Figure 3F, Supplemental Figure 5). This suggests that loss of Heh1 increases CNV in an unbiased fashion, while selection acts to determine if the *ENA* copy number expands or contracts depending on fitness for the environment.

To understand how *HEH1* and *HEH2* achieve this effect, we investigated their synthetic genetic interaction networks [55-60]. Interestingly, *HEH1* shows synthetic sickness with genes involved at different stages of transcription: *TOP1*, which prevents negative DNA supercoiling, the THO-TREX/TREX2 complex, which influence transcriptional elongation and termination (coordinated with mRNA export), and *XRN1* and *RRP6*, which participate in RNA surveillance and degradation. All these genes play a major role in preventing the formation or persistence of RNA-DNA hybrids (R-loops) that constitute a major threat to genome stability [61]. Intriguingly, *HEH2* consistently displays the opposite effect of *HEH1*, acting as a genetic suppressor of the same genetic network. For example, deletion of *HEH2* substantially rescues the growth of synthetically sick *heh1*Δ*sac3*Δ, *heh1*Δ*xrn1*Δ and *heh1*Δ*rrp6*Δ strains (Figure 4B, Supplemental Figure 6). Interestingly, the deletion of the Heh2 winged-helix domain is enough to partially rescue the fitness loss of *heh1*Δ*sac3*Δ (Figure 4B). These genetic interactions suggest the possibility that Heh1 and Heh2 might modulate R-loop formation or resolution, which we first tested genetically by examining interactions between *HEH1* and *HEH2* and the ribonuclease H1 and H2 enzymes involved in the removal R-loops in *S. cerevisiae*, encoded by *RNH1* and *RNH201*[62]. Loss of *HEH2* is suppressive of the growth defect of the *rnh1*Δ*rnh201*Δ genotype in the presence of hydroxyurea, driving a marked increase in fitness with the colony size increasing by ≈75% (Figure 4C).

**Figure 4.**
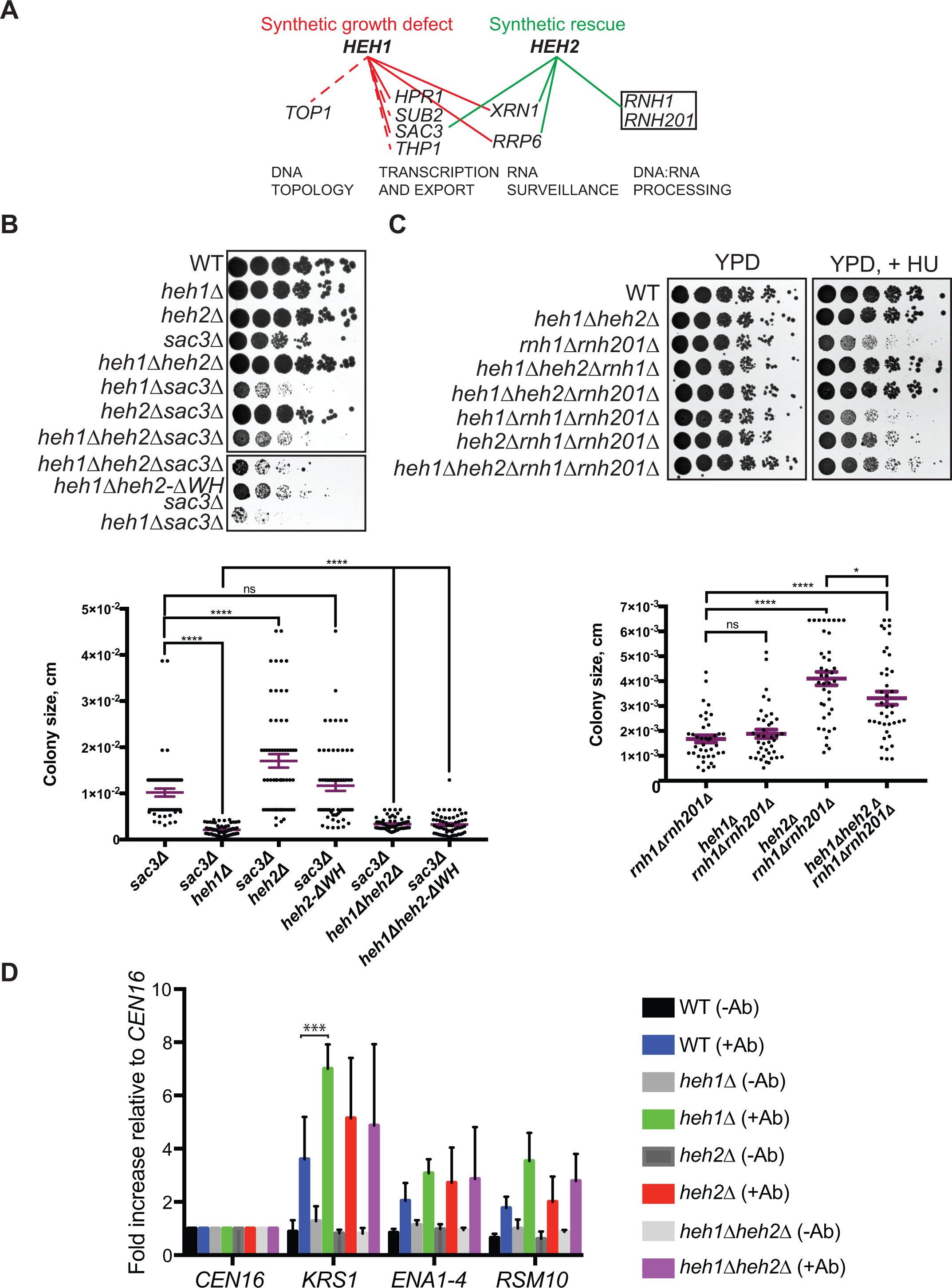
*HEH1* and *HEH2* are part of a genetic pathway modulating R-loop formation. (A) Schematic of the tested genetic interactions (red and green lines) and those reported in the literature (red dashed lines). Red lines indicate a synthetic growth defect and green lines indicate a synthetic growth rescue for the connected gene pairs. (B) Synthetic genetic interactions between *SAC3* and *HEH1/2.* Serial dilutions, 1:10, cells were grown on YPD plates. Quantification of the colony size (expressed in cm^2^), for 60 colonies for each strain from three independent experiments are plotted. Mean ± SEM.^****^ *p* < 0.0001 (ANOVA). (C) Synthetic genetic interactions between *RNH1/201* and *HEH1/2.* Serial dilutions, 1:10, cells were grown on YPD and on YPD containing 50 mM HU. Measurements of the colony size (expressed in cm^2^) for 60 colonies for each strain from three independent experiments are plotted. Mean ± SEM. **** *p* < 0.0001 (ANOVA). n.s. is not significant. (D) R-loop analysis by DRIP-qPCR across the *ENA* genomic locus in*WT*, *heh1*Δ, *heh2*Δ, *heh1*Δ*heh2*Δ strains. *CEN16*, *KRS1, ENA1-4,* and *RSM10* were analyzed by qPCR. The values for no-antibody (-Ab) and antibody S9.6 (+Ab) were calculated as described in Methods, and normalized to *CEN16* to compensate for differences in IP efficiency. Mean ± SD of three biological replicates, each analyzed by qPCR three times.

To directly test how *HEH1* and *HEH2* influence the accumulation of R-loops at the *ENA* locus, we carried out targeted RNA-DNA immunoprecipitation (DRIP) using the monoclonal antibody S9.6 [63] followed by qPCR. We detected robust RNA-DNA hybrids across the *ENA* region (Figure 4D). However, there is a marked asymmetry, with the gene *KRS1* accumulating almost twice the extent of hybrids detected at the *ENA* or *RSM10* genes, thus correlating with the stability of the exogenous *URA3* gene inserted at these locations (Figure 2). Interestingly, the deletion of *HEH1* drives an overall increase of RNA-DNA hybrids, most dramatically at *KRS1* (Figure 4D); *heh2Δ,* or the combined*heh1*Δ *heh2*Δ genetic backgrounds do not produce any measurable change compared to WT cells, suggesting that loss of *HEH1* only drives accumulation of RNA-DNA hybrids when *HEH2* is present. Taken together, these orthogonal approaches suggest that the presence of *HEH2* favors the formation RNA-DNA hybrids, while *HEH1* opposes this activity.

## Discussion

This work provides evidence for pathway in which the act of transcription and the LEM domain protein *HEH2* promote CNV at the *ENA* locus, while *HEH1* opposes this activity. In cells lacking *HEH1*, RNA-DNA hybrids accumulate downstream of the *ENA* locus, which is associated with a more rapid rate of both CNV expansion (in the presence of selective pressure) and contraction (in the absence of selective pressure). This model is further supported by a network of opposing genetic interactions between *HEH1* and *HEH2* and genes encoding factors influencing transcription termination, RNA turnover, or metabolizing RNA-DNA hybrids. We speculate that the expansion of the *ENA* locus and salt tolerance provides just one example of the biological contexts in which specific sites in the genome undergo this type of stress-induced CNV pathway, although further studies will be required test this notion.

While it has previously been appreciated that tandem genomic repeats frequently undergo reciprocal exchanges that can drive changes in copy number [38] (which can then be acted upon by selection) and that accumulation of RNA-DNA hybrids is tied to local, increased genome instability [61], these processes are largely thought to take place due to faulty repair mechanisms or incidental (and often deleterious) effects. At the same time, substantial evidence supports the notion that CNV is exploited to promote adaptation in yeasts [64, 65], but also in a broad array of other organisms and context, for example resistance to antibiotics in prokaryotes or chemotherapy resistance in the setting of cancer (reviewed in [7, 66]). Here we provide evidence that the antagonistic functions of conserved LEM domain proteins [67], likely through modulating RNA-DNA hybrids, influence the local loss of “genome stability” and fuel CNV-dependent adaptation. Our findings extend previous recent observations that CNVs can be driven in response to environmental stress in a manner that requires both transcription [2] and its associated histone modifications [2, 68]. Importantly, here we find that such processes result not only from stochastic losses of genome integrity resulting from collisions between the replication fork and RNA-DNA hybrids and subsequent repair processes, but also from a balance of *HEH1*- and *HEH2*-dependent influences that regulate a local loss of genome stability in response to physiological cues. Importantly, in this context we conclude that spontaneous gain/loss of copy number and selection for copy gain (growth in LiCl) alone cannot explain our observations.

How might the balance of LEM domain protein activities influence RNA-DNA hybrid formation? It is interesting to note that paralogues (in this case the *HEH1*and *HEH2* ohnologues arising from the whole genome duplication in the history of *S. cerevisiae*) often evolve to carry out antagonistic functions [69, 70]. The genetic interaction network of *HEH1*and *HEH2* (Fig. 4) supports the notion that these factors antagonize each other to regulate either transcriptional termination, leading to stalled RNA-DNA hybrids, or instead the removal of RNA-DNA hybrids by RNase H. Interestingly, the domain of Heh2 necessary to impart its activity, its C-terminal winged helix domain, is structurally shared among the RNA-interacting components of the THO-TREX complex [71], although further studies will be necessary to define the biochemical activities or interactions by which these LEM domain proteins influence the accumulation of RNA DNA hybrids.

In addition to changes in copy number, here we also demonstrate that the ENA locus is also a hot spot for point mutations, in a manner that is at least partly influenced by its transcriptional response to high concentrations of LiCl. For example, the rates of mutation within *URA3* 5’ and 3’ to the *ENA* gene cluster is substantially higher than the median rate observed across tens of *URA3* insertion sites on chromosome VI [40], raising the question of why this locus is so unusually unstable. As the *ENA* locus is replicated early [72], and the occupancy of Pol II is similar both up and downstream of the ENA locus [73], we favor the idea that, as was described for the acquisition of suppressor of ochre mutations in budding yeast due to non-random mutations to one of the six tRNA-Tyr genes [74], the orientation of the *ENA* transcriptional unit (and the associated RNA-DNA hybrids) drives collisions with the replication fork that initiates from a nearby ARS, only 21 kbps 5’ to the *ENA* gene cluster. The change of the mutational signature in the presence of LiCl downstream of the *ENA* locus is compatible with the mutation profile observed for DNA polymerases involved in gene conversion (Pol and Pol) interrogated at another highly unstable region of the budding yeast genome, the mating type locus [75], suggesting that they may be products of DNA repair acting after replication fork collapse, although translesion synthesis, which can drive single base pair substitutions, may also play a role [76-78]. This model echoes the observations that the recombinase RecA and DNA synthesis are required for “directed mutagenesis” in bacteria [79, 80] but supports the existence of mechanisms that drive a “regional” (rather than random) mutagenesis followed by selection rather than a preference for adaptive mutations.

Oncogene amplification represents a major driver of carcinogenesis and a challenge to cancer therapy [81-83]. The results presented here suggest that adaptive mechanisms that drive CNV can be disabled to combat oncogenesis and therapy resistance. The antagonistic effect of *HEH1* and *HEH2* on this process highlights the need for further investigation into how members of the LEM domain family in mammals, which includes LAP2, emerin, LEM2 and MAN1, impact on genome integrity; to date this potential connection has gone largely uninvestigated [84].

## Materials and Methods

### Yeast strain generation and culturing

All yeast strains used in this study and their derivation are listed in Table S2. Unless otherwise stated, all experiments were conducted at 30°C. The *rpb1-1* strain (derived from KWY1302, a gift K. Weis, ETH Zurich, Switzerland) was cultured at room temperature (RT) as described in the Figure Legends. All strains were grown in YP (1% yeast extract and 2% peptone) with 2% dextrose (YPD) with the addition of different concentration of LiCl, as described. Standard yeast manipulations including transformations, tetrad dissection, and PCR-based integration were performed as described[85]. The LacO integrations were generated as described[86].

### *In vitro* evolution experiment

The strains were initially grown overnight at 30°C in 2 ml YPD. In the morning, the density of the culture was quantified using an automated cell counter (Moxi^z^- Orflo) and 50,000 cells were diluted into 50 ml of YPD with100mM LiCl and grown for 24h. This procedure of quantification and dilution of the culture was repeated for 18 days (~200 generations). Every day samples were collected and stored for fitness and gene copy number analysis.

### Reverse transcription PCR and qPCR

RNA was prepared using the MasterPure Yeast RNA purification Kit (Epicentre) according to manufacturer’s instructions from cells growing in exponential phase at 30°C in YPD, and after 10 min exposure to YPD with 100mM LiCl. DNA contamination was removed by treating samples with DNaseI for 45 min at 37°C. cDNA was synthesized from 500ng of total RNA using Superscript^®^ III First-Strand Synthesis (Invitrogen) with oligo-dT primers. cDNA was added to the iTaq™ Universal SYBR^®^ Green supermix (BIORAD) with primers to amplify *ENA1-4* (the primers anneal in regions that are identical between all 4 gene copies) and *ACT1* as internal load control. Reactions mixes were cycled in a CFX96 Touch Real-Time PCR Detection system (BIORAD). To calculate relative gene expression we used the 2-ΔΔCt method of analysis[87]. Log2 2^-ΔΔCt^ values were from three independent experiments were normalized to control condition (0mM LiCl) and plotted as shown in Figs. 1B, 3E. Primers listed in Table S3.

### Genomic DNA extraction and copy number quantification by qPCR

Genomic DNA was prepared using a modified Winston method. Cells were grown overnight to saturation in YPD, washed with 1 ml of water and resuspended in 10 mM Tris-HCl, pH 8.0, 2% TRITON X-100, 1% SDS, 100 mM NaCl, 1 mM EDTA. 200 µl of phenol:chloroform:isoamyl alchol (25:24:1) (Fisher Scientific) and 100 µl of glassbeads were added to the cell suspension and vortexed for 5 min. After adding 200 µl of 10 mM Tris-HCl pH 8.0, 1 mM EDTA (TE), the suspension was centrifuged for 5 min at 17000 x g, the supernatant transferred to a new tube and genomic DNA was precipitated with 100% ethanol. The pellets were washed with 70% ethanol, air dried and resuspended in TE containing 50 µg/µl RNase. The method adopted for measuring copy number was adapted from Weaver et al.[88] with minor modifications. Briefly, we used the following approach: 2^-ΔΔCq^ based on a target assay T (*ENA*) for the DNA segment being interrogated for copy number variation and a reference assay R (*ALG9*) for an internal control segment which is a single copy gene. The Δ*C*_*q*_ = (*C*_*q*,*ENA*_-*C*_*q*,*ALG9*_) is a measure of the copy number of the target segment (*ENA*) relative to the reference segment (ALG9). The next step in determining the relative copy number is to calibrate the Δ*C*_*q*_ value to a sample with a single copy number for the target (*ENA*) and for reference gene (*ALG9*) Δ*C_q,C_*= (*C_q,ENA_*-*C_q,ALG9_*). Assuming that the efficiencies of the target and reference assay are similar and close to 1[87], the relative copy number is calculated from the formula 2-^ΔΔ*Cq*^ where ΔΔ*C_q_* = Δ*C_q_* - Δ*C_q,c_*. For every target and calibrator sample, three different concentrations of genomic DNA were tested, each in triplicate, thus allowing us to evaluate the efficiency for every sample analyzed.

### Genome stability assay

Cells were grown in selective media (CSM-URA) for two days starting from a single colony obtained from a freshly re-streaked strain. After two days, the cells were resuspended in complete YPD media for a single cell cycle (2 hrs. for WT strains) and then plated on YPD plates (to determine the total number of cells plated) and on 5-fluoorotic acid (5-FOA)-containing plates to identify the number of cells that lost URA3 activity. In the case of LiCl treatment, cells were released for a single cell cycle (2 hrs. for WT) in YPD containing 100 mM LiCl. The data are reported as a ratio between the number of cells growing in 5-FOA plates versus the total number of cells measured on YPD plates.

### Microscopy

For the imaging experiments, cells were grown to mid-log phase and immobilized on a 1.4% agarose pad containing complete synthetic medium (CSM) with 2% glucose and sealed with VALAP (1:1:1 Vaseline/lanolin/paraffin). The microscopy experiments were carried out on a wide-field deconvolution microscope (DeltaVision; Applied Precision/GE Healthcare) equipped with a 100×, 1.40 NA objective lens and solid state illumination. The images were acquired using an Evolve EMCCD camera (Photometrics). Temperature control was achieved through the enclosure of the microscope within an environmental chamber. In all cases, a z-series of images with 200 nm spacing were acquired and further processed as described under “Image processing and analysis.”

### Image processing and analysis

The 3D reconstruction of the nuclear envelope, fitting of the LacI-GFP/LacO focus and the position of the LacI-GFP/LacO with respect to the nuclear envelope were determined as described in our published work[89].

### Single nucleotide polymorphism analysis

Variant data was obtained for 21 *S. cerevisiae* strains, kindly provided by Dr. J. Michael Cherry[49], which includes BC187, BY4741, BY4742, CEN.PK2-1Ca, D273-10B, DBVPG6044, FL100, FY1679, JK9-3d, K11, L1528, RedStar, RM11-1A, SEY6210, Sigma1278b-10560-6B, SK1, UWOPS05_217_3, W303, X2180-1A, YPS163, and YS9. The genome build used as the reference and for plots was UCSC sacCer3. Individual SNPs were excluded if any of the following criteria were met: allele balance for heterozygous genotype > 0.75, read depth > 360, strand bias > −0.1, or those flagged for low quality. These 21 filtered files were combined and a variant frequency file was generated, which lists the number of strains for which a variant was identified at each position for each chromosome. The variant frequency file was divided by the total number of variants across the genome (729,305) to make a distribution file. Peaks were called (and island scores were determined) using SICER[90] with the following settings: window size = 100 bp, gap size = 100 bp, E-value = 100, FDR of 2.1%.

### Epistasis analysis

Genetic interactions were assessed by spotting equivalent numbers of cells in six 10-fold serial dilutions incubated at 30°C for 2-4 days. For epistasis analysis the size of 60 colonies from three independent growth assays were measured using Fiji[91]. For each replicate the strains tested were isolated directly from spores obtained from tetrad dissection.

### DNA:RNA immunoprecipitation (DRIP)

Yeast strains grown in YPD were harvested in exponential phase, washed with 1 ml of water and frozen in liquid nitrogen. The cells were lysed with RA1 buffer (Macherey-Nagel) supplemented with 100 mM NaCl and 1% i-mercapthoethanol. The cell suspension was mixed with phenol:chloroform:isoamyl alcohol (25:24:1) (Fisher Scientific) and glass beads. Cells were broken by mechanical shaking using a pulsing vortex mixer. After spinning, the upper phase was transferred to a new tube and nucleic acids were precipitated with 1 ml of isopropanol followed by centrifugation. Pellets were washed with 70% ethanol, air dried, resuspended in 50 mM Tris-HCl, 75 mM KCl, 3 mM MgCl_2_, 10 mM DTT, 5 mM EDTA and sonicated with a Bioruptor (Diagenode) to obtain 100-500 bp fragments. Fifteen micrograms of sonicated nucleic acids were diluted in 1 ml of IP buffer (0.1% SDS, 1% Triton X-100, 10 mM HEPES pH 7.7, 0.1% sodium deoxycholate, 275 mM NaCl) and incubated overnight on a rotating wheel at 4°C with 1 Cg of S9.6 antibody (Kerafast), followed by precipitation with 25 µl of Dynabeads protein G (Life Technologies) pre-blocked with bovine serum albumin and *E. coli* DNA. Beads were washed 5 times with IP buffer, and the DNA fragments were collected by PCR cleanup kit (QIAGEN). The collected DNA fragments were quantified by qPCR using the primers listed in Table S3, and values for DRIP were calculated using the formula Δ*C_q_* “- antibody”= 2^-(C*q* “beads only” – C*q* “input chromatin”)^ and Δ*C_q_* “+ antibody”= 2^-(C*q* “beads only” – C*q* “input chromatin”)^. The values were normalized to *CEN16*, which was set to a value of 1 in order to compensate for differences in immunoprecipitation efficiency[92].

## Colombi et al., Supplemental Figures 1-6, Supplemental Tables 1-3

**Supplemental Figure 1.**
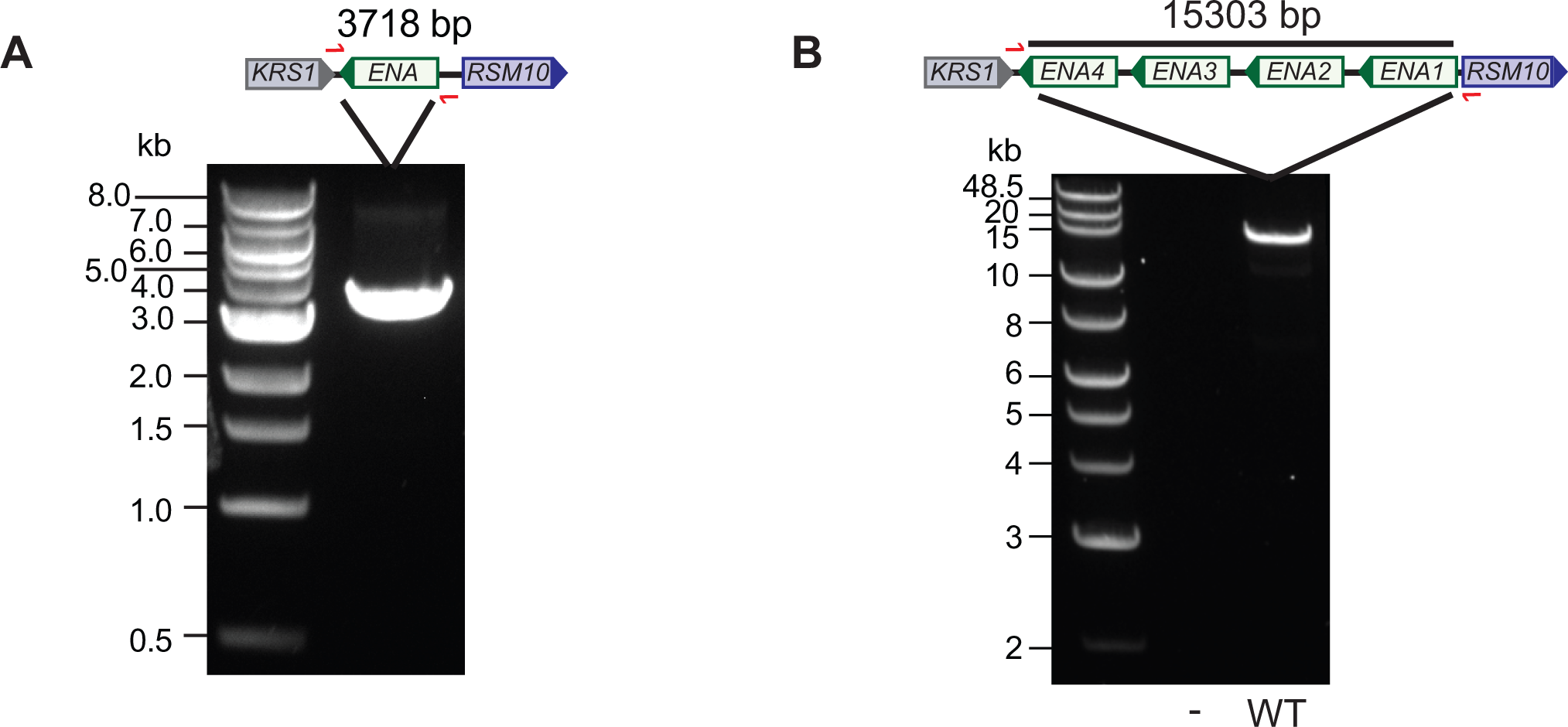
Characterization of *ENA* genomic locus in W303. (A) PCR amplification of the *ENA* genomic locus from genomic DNA extracted from a strain containing a single *ENA* gene. The locus is schematized above the gel and the red arrows indicate the position of the PCR primers. (B) PCR amplification of the *ENA* genomic locus of our WT W303 strain finds four *ENA* gene copies (validated by qPCR, Fig. 1F).

**Supplemental Figure 2.**
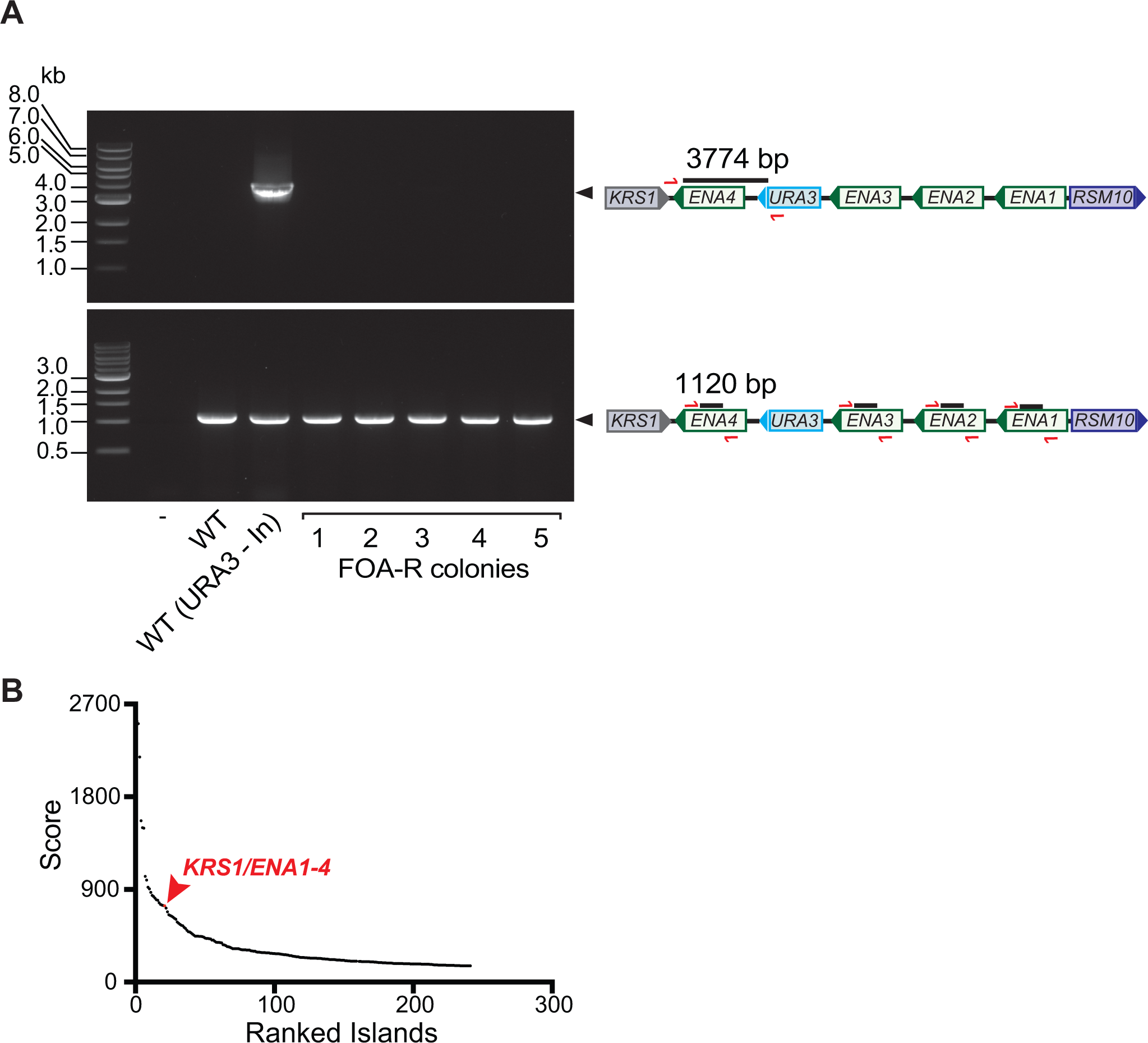
(A) The *URA3* gene inserted between the *ENA* repeats (“In”) is lost in the clones that become resistant (FOA-R) to growth on 5-FOA. 1 to 5 represents five different clones analyzed. The PCR in the lower panel shows the presence of the *ENA* genes even in clones that lost *URA3*. On the right are the schematics of the PCR primer binding sites (red) at the starting locus prior to 5-FOA selection. (B) The genomic region 3’ to the *ENA* locus has a very high SNP load as assessed by comparative genomics (red, see also Table 1).

**Supplemental Figure 3.**
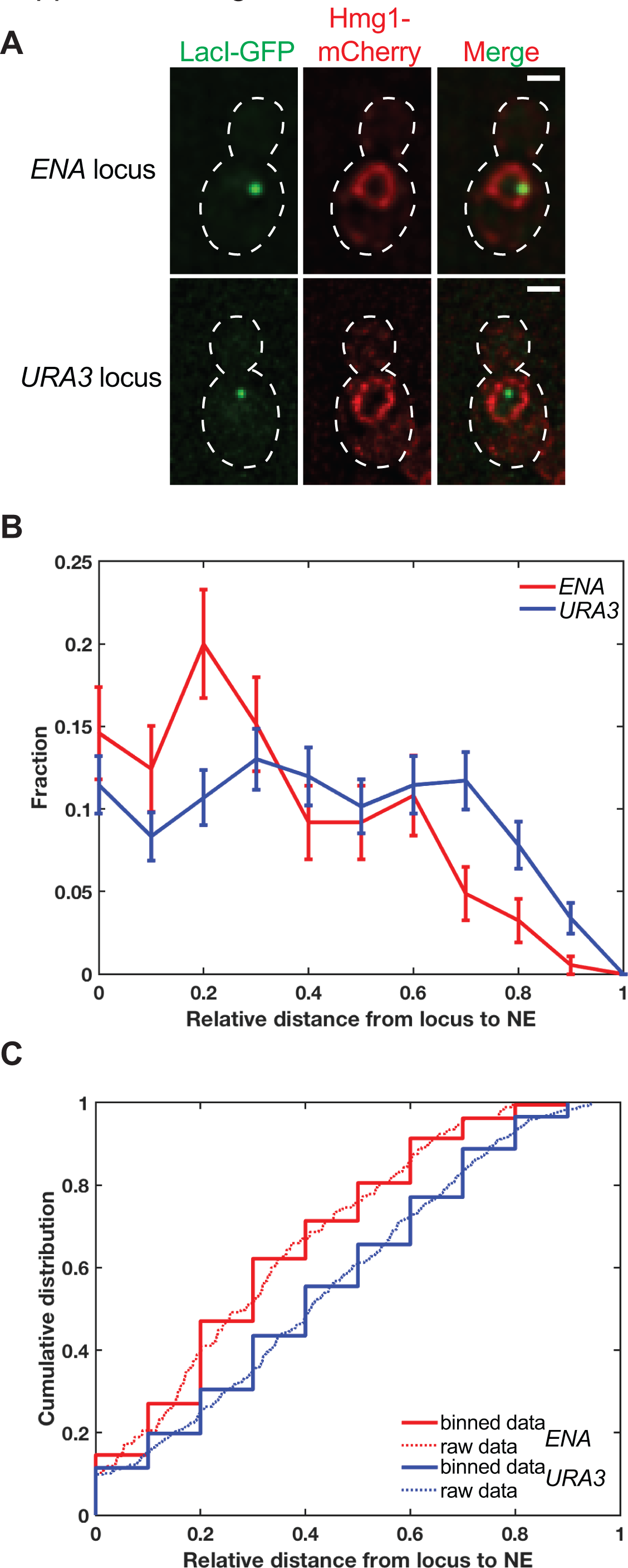
The *ENA* locus is associated with the nuclear periphery. (A) Fluorescent micrographs of one *z* section of cells expressing Hmg1-mCherry and LacI-GFP. The Lac operator (LacO) array was inserted in proximity to the *ENA* locus. Green and red channels are shown, in addition to the merge. Dotted lines denote cell boundaries. (B) Distribution of the normalized distance between the LacO array and the nuclear envelope (NE) derived from the 3D reconstructions. Here the *ENA* locus is enriched at the NE and it is depleted the nuclear interior, while the endogenous *URA3* locus is relatively depleted from the NE and enriched at the nuclear interior. (C) Cumulative distribution of the normalized distance of the LacO array from the NE. The dashed lines represent the raw data while the continuous lines represent the binned data.

**Supplemental Figure 4.**
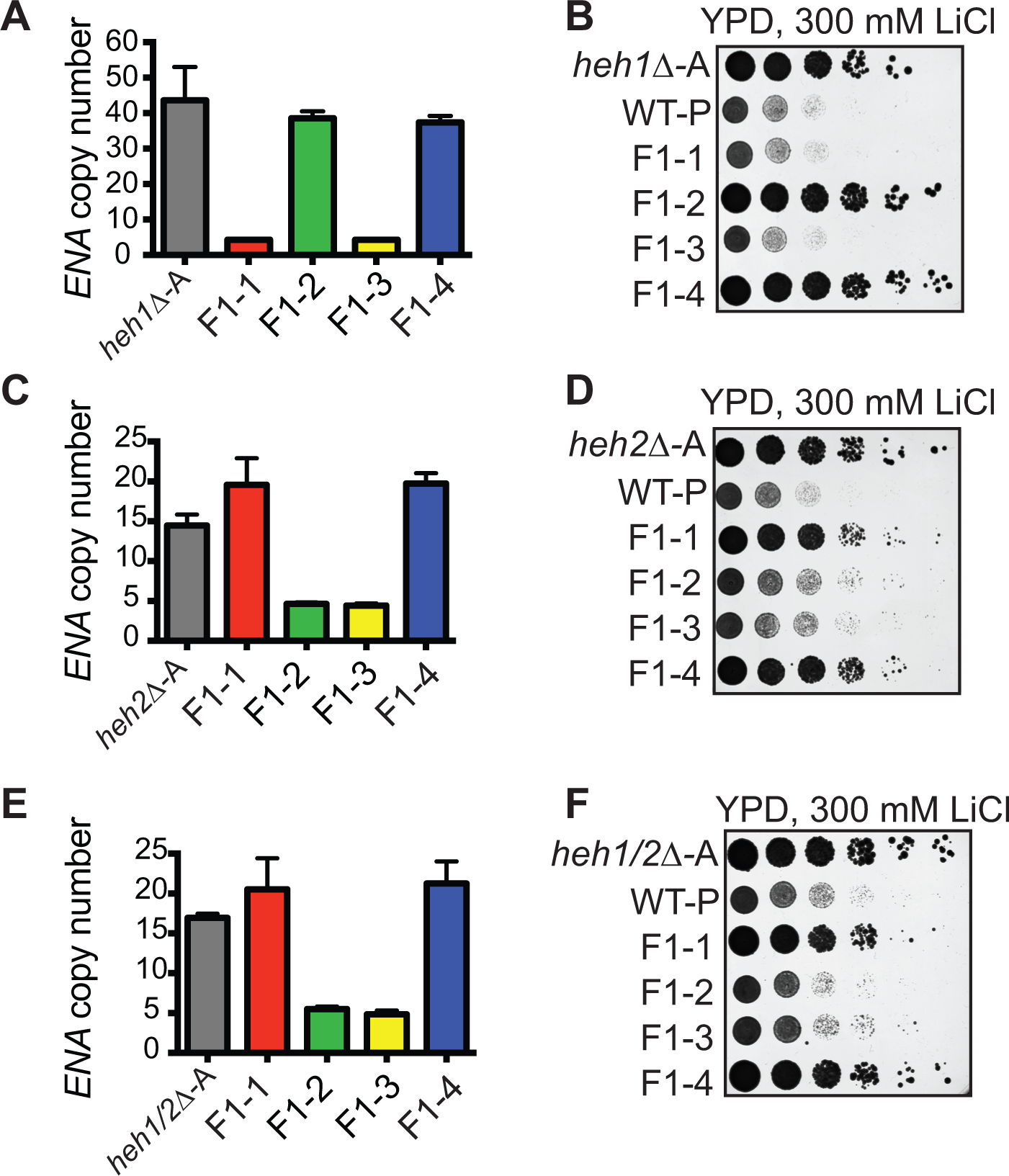
The *ENA* locus undergoes intra-chromosomal expansion in *heh1*Δ, *heh2*Δ and *heh1*Δ*heh2*Δ cells. (A, C, and E) The adapted *ENA* locus segregates in a Mendelian fashion in the adapted (A) *heh1*Δ, *heh2*Δ and *heh1*Δ*heh2*Δ strains, respectively. The *ENA* gene copy number of the four spores (F1-1-F1-4) was assessed by qPCR. (B, D, F) The spores from (A, C and E) that inherit the adapted *ENA* locus also show improved fitness on media containing 300 mM LiCl over those inheriting the WT *ENA* locus. Serial dilutions as in Fig. 1C.

**Supplemental Figure 5.**
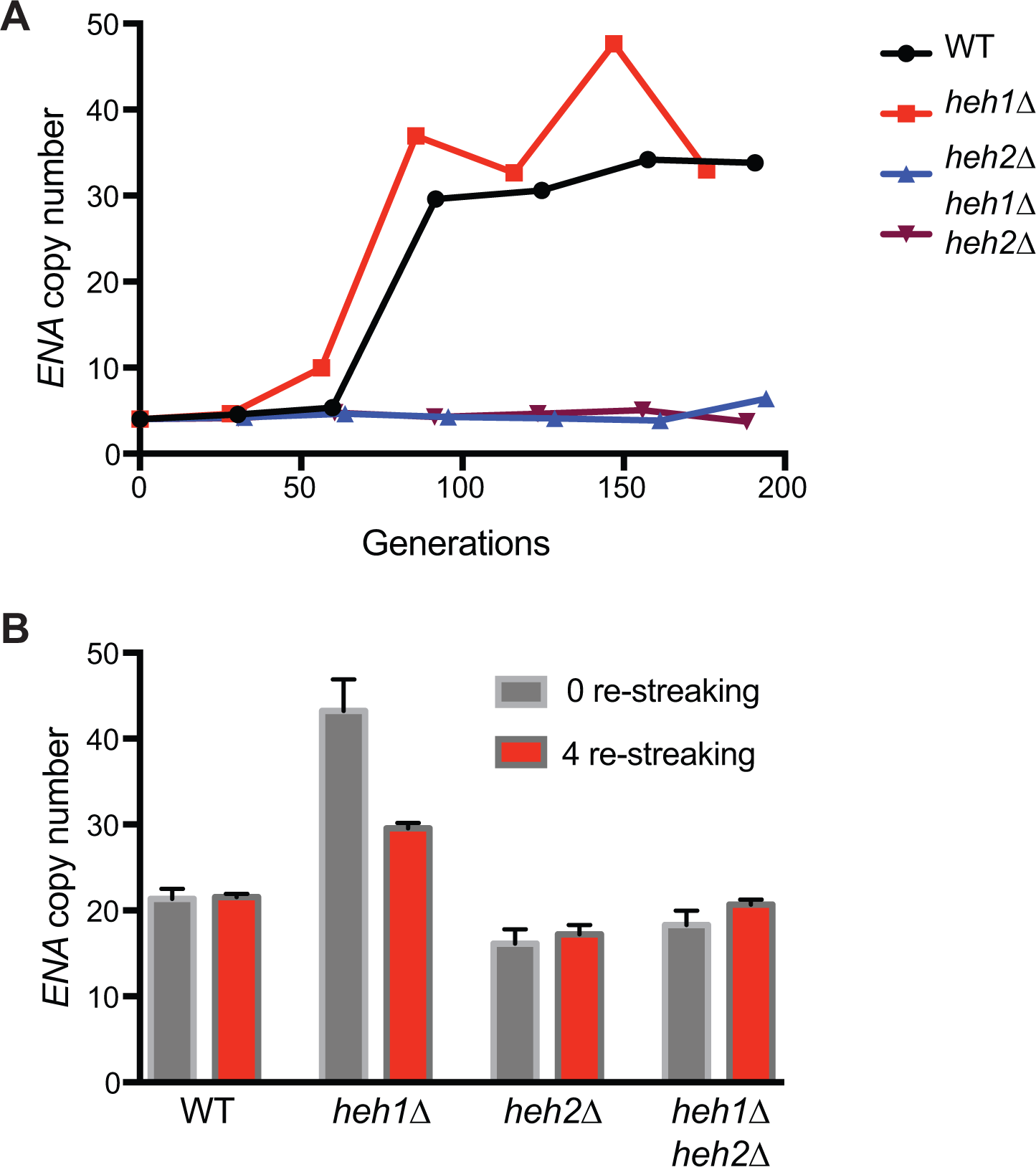
Independent biological replicates of *ENA* expansion and contraction. (A) qPCR to determine *ENA* copy number through the *in vitro* evolution experiment of the indicated strains. (B) Second independent experiment showing that *HEH1* influences the stability of the adapted *ENA* locus in the absence of selective pressure. qPCR analysis of *ENA* copy number of adapted strains before and after 4 serial re-streakings on complete media.

**Supplemental Figure 6.**
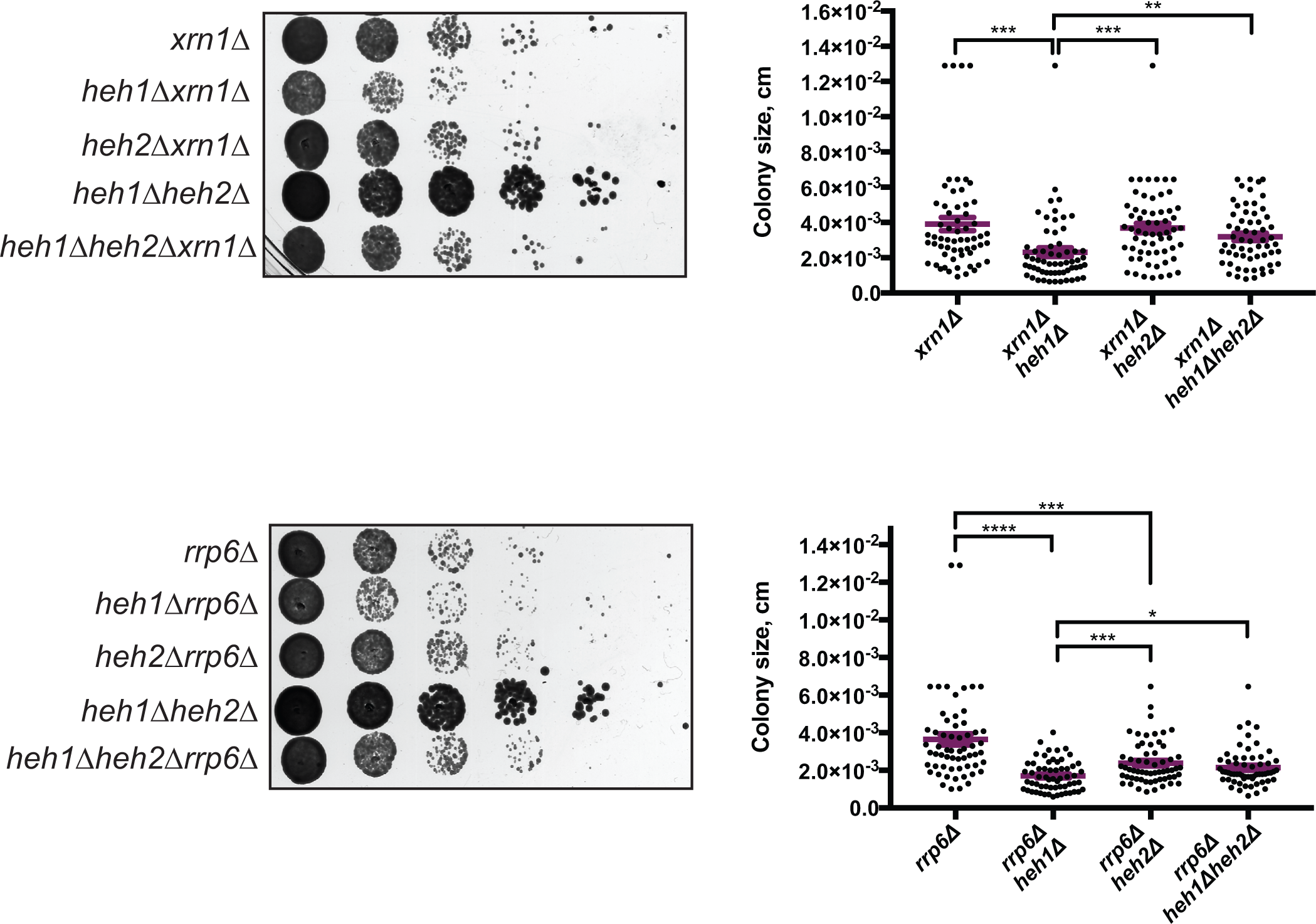
Synthetic genetic interactions between *XRN1* (top), *RRP6* (bottom) and *HEH1/2.* Serial dilutions as in Fig. 1C. Measurement of the colony size (expressed in cm^2^); 60 colonies from each strain from three independent experiments are plotted. Mean ± SEM. ^***^ *p* < 0.001, ^**^ *p* < 0.01 (ANOVA).

### Supplemental Tables

**Table 1.**
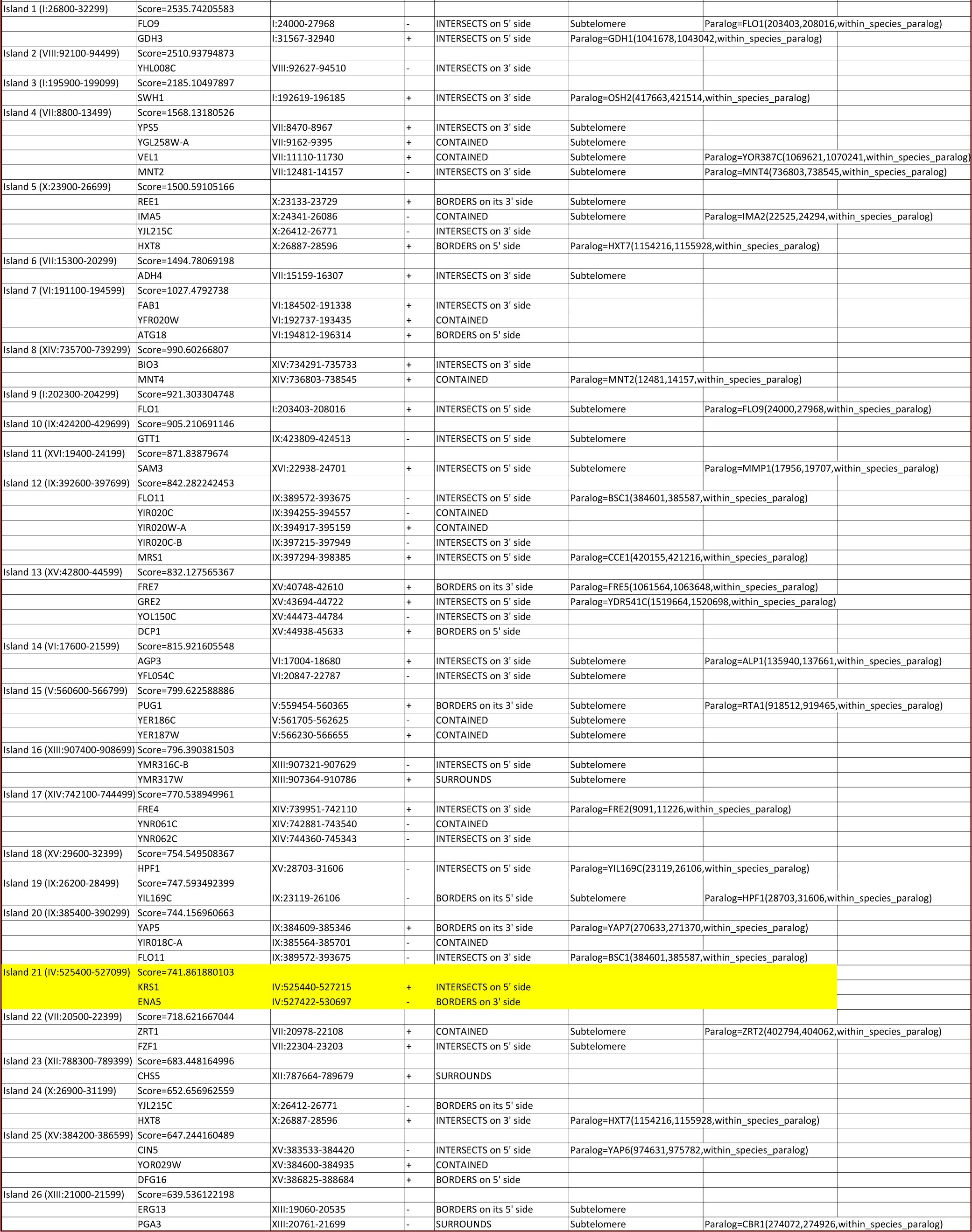
Top SNP islands (see Methods). Coordinates for each island, as well as gene features, whether the region is in the subtelomere (defined as within 25 kb of the chromosome end) and whether the region contains a gene with a paralogue are indicated. The *KRS1/ENA* region, ranked 21^st^, is highlighted.

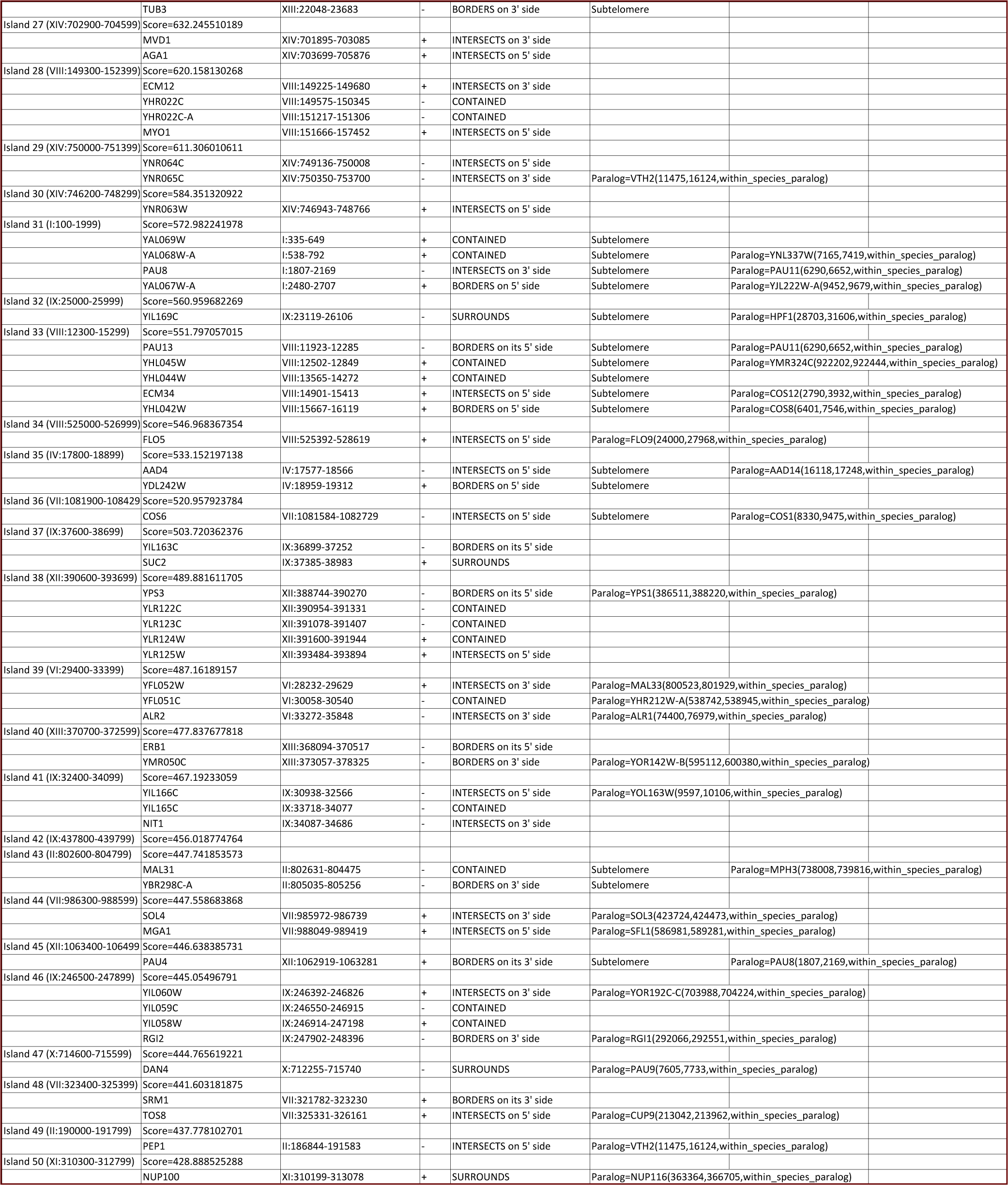

**Table 2.**
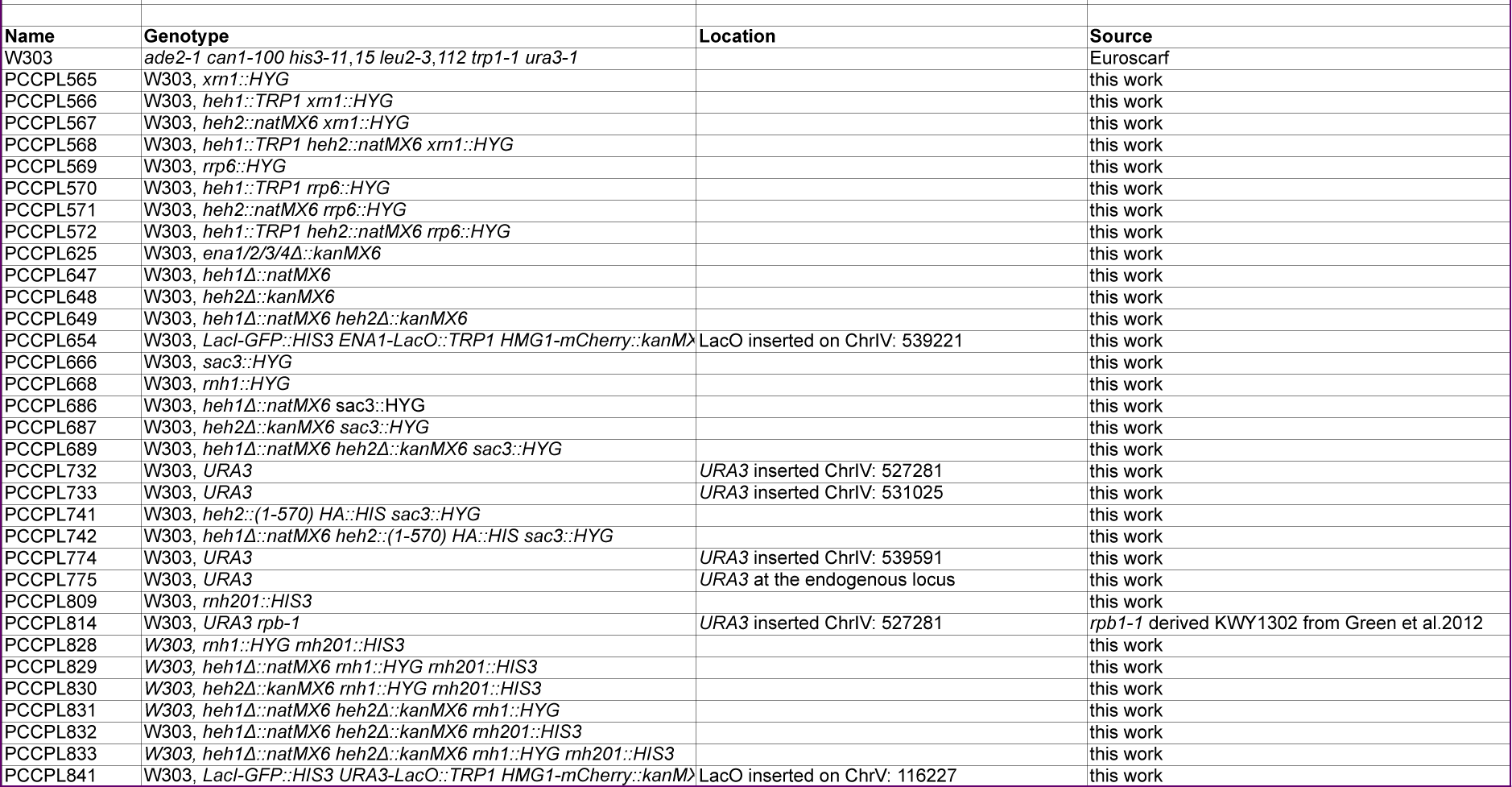
*S. cerevisiae* strains used in this study.

**Table 3.**
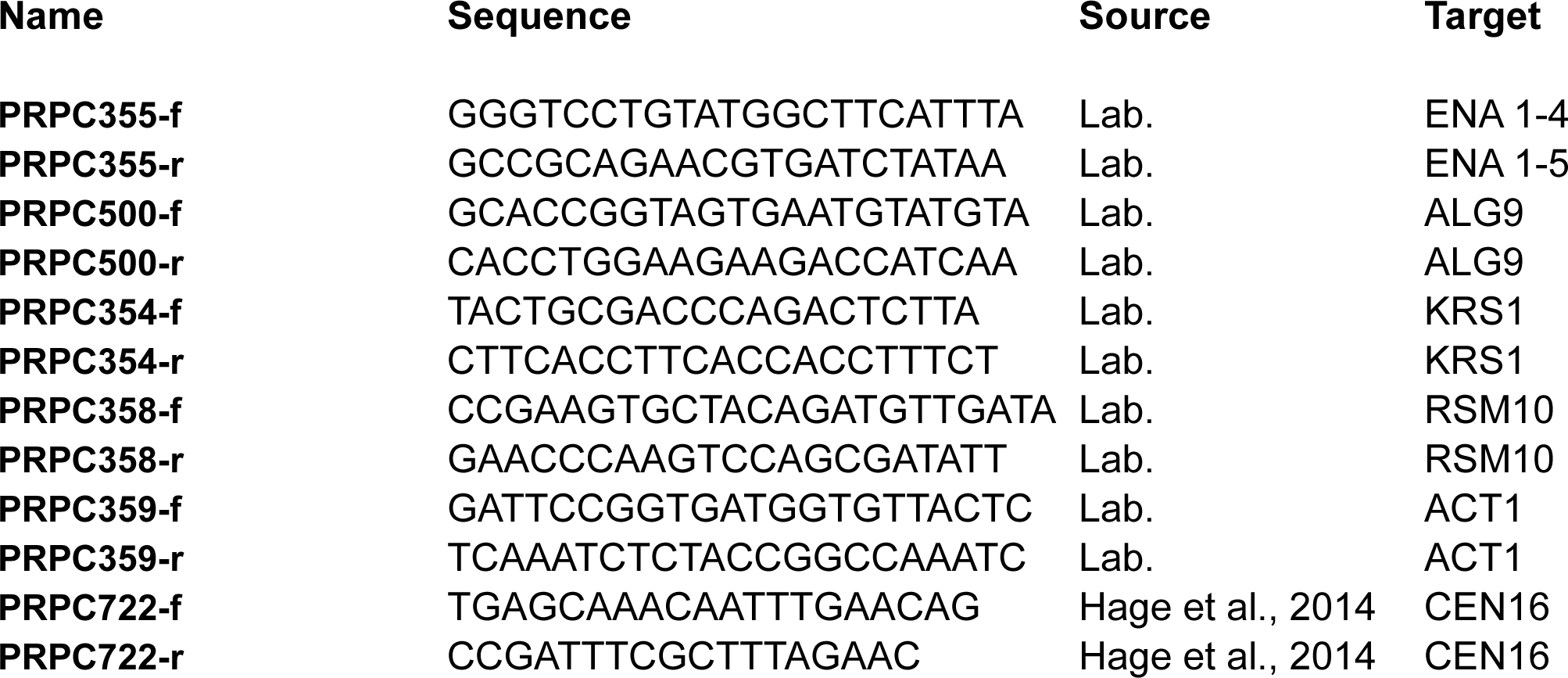
Oligonucleotide primers used in this study.

